# Multimodal system for recording individual-level behaviors in songbird groups

**DOI:** 10.1101/2022.09.23.509166

**Authors:** L. Rüttimann, Y. Wang, J. Rychen, T. Tomka, H. Hörster, R.H.R. Hahnloser

## Abstract

The implicit goal of longitudinal observations of animal groups is to identify individuals and to reliably detect their behaviors, including their vocalizations. Yet, to segment fast behaviors and to extract individual vocalizations from sound mixtures remain challenging problems. Promising approaches are multimodal systems that record behaviors with multiple cameras, microphones, and animal-borne wireless sensors. The instrumentation of these systems must be optimized for multimodal signal integration, which is an overlooked steppingstone to successful behavioral tracking.

We designed a modular system (BirdPark) for simultaneously recording small animals wearing custom low-power frequency-modulated radio transmitters. Our custom software-defined radio receiver makes use of a multi-antenna demodulation technique that eliminates data losses due to radio signal fading and that increases the signal-to-noise ratio of the received radio signals by 6.5 dB compared to best single-antenna approaches. Digital acquisition relies on a single clock, allowing us to exploit cross-modal redundancies for dissecting rapid behaviors on time scales well below the video frame period, which we demonstrate by reconstructing the wing stroke phases of free-flying songbirds. By separating the vocalizations among up to eight vocally interacting birds, our work paves the way for dissecting complex social behaviors.

## Introduction

Acoustic communication is vital for many social behaviors. However, to study interactions among animals that are kept in groups entails many measurement challenges beyond the already considerable challenges of analyzing longitudinal data of isolated animals (Lipkind et al. 2017; Tchernichovski et al. 2001; Kollmorgen, Hahnloser, and Mante 2020). One of the key difficulties of group-level behavior research is to perform automatic recognition of individuals and their actions. Action recognition has emerged as the computational task of detecting behaviors from video or audio, or from signals collected with animal-borne sensors.

In animal behavior studies, video recording stands as the quintessential and most prevalent method for monitoring behaviors. Video-based action recognition has traditionally been based on posture tracking (Segalin et al. 2020; Fujimori, Ishikawa, and Watanabe 2020; Perkes et al. 2021) to avoid data-hungry training of classifiers on high-dimensional video data.

Recently, posture tracking has greatly improved thanks to deep-learning approaches (Nath et al. 2019; Mathis et al. 2020; Walter and Couzin 2021; Badger et al. 2020; Harley, Fang, and Fragkiadaki 2022; Doersch et al. 2023; Naik et al. 2023; Pereira et al. 2022). Action recognition from video requires good visibility of focal animals because visual obstructions tend to hamper recognition accuracy. Given that freely moving animals may occlude one another, e.g., in birds during nesting, there seems to be a limitation to the usefulness of pure vision-based approaches.

Sound recordings have also been instrumental in action recognition. Sounds can be informative about both vocal and non-vocal behaviors. For example, wing flapping during flying produces a characteristic sound signature. But also preening, walking, and shaking can be recognized from sounds (Stowell, Benetos, and Gill 2017). The task of classifying sounds is known as acoustic scene classification (Barchiesi et al. 2015). Microphones record not just the focal animal but also background sounds, which makes sound-based action recognition and actor identification challenging tasks, especially when many animals interact with one another.

Microphone arrays can improve localization and separation of sound sources (Rhinehart et al. 2020), though with arrays alone it remains difficult to assign vocalizations to individuals. Further improvements are obtained with multimodal approaches combining cameras and microphone arrays, for example to assign vocalizations to individual mice in a social home cage (Heckman et al. 2017) or in dairy cattle in a barn (Meen et al. 2015). Many multimodal systems for action recognition make use of motion-tracking devices, for example to quantify gesture–speech synchrony in humans (Pouw, Trujillo, and Dixon 2020).

Methods that integrate different data modalities have been deemed particularly promising for functional decoding of animal communication (Rutz et al. 2023). One challenge with multimodal systems is the synchronization of the various data streams. Usually, each sensor modality is recorded with a dedicated device that uses its own internal sampling clock, and these clocks tend to drift apart. Furthermore, the recordings must be started at exactly at the same time on all devices, or the individual data streams must be aligned post-recording using markers in the sensor signals or auxiliary synchronization channels, which are labor intensive and error-prone processes (Pouw, Trujillo, and Dixon 2020; Zimmerman et al. 2009; Dolmans et al. 2021).

In general, limitations due to sight occlusions and sound superpositions can be overcome with animal-borne sensors such as accelerometers (Anisimov et al. 2014; Eisenring et al. 2022)gyroscopes, microphones (Ter Maat et al. 2014) and global positioning systems (GPS) (Eisenring et al. 2022). In combination with wireless transmitters (Ter Maat et al. 2014) and loggers (Anisimov et al. 2014), these sensors enable the detection of behaviors such as walking, grooming, eating, drinking, and flying, for example, in birds (Gómez Laich et al. 2009), cats (Watanabe et al. 2005), and dogs (Gerencsér et al. 2013), though often with low reliability due to noisy and ambiguous sensor signals (Stowell, Benetos, and Gill 2017).

One drawback of animal-borne sensors is that they can impact behavior. In zebra finches their effects on locomotion and singing rate are temporary: right after attachment of the sensors, the singing rate and amount of locomotion both tend to decrease but return to baseline within 3 days when sensors weigh 0.75 g (Gill et al. 2016) and within 2 weeks when sensors weigh 3 g (Anisimov et al. 2014). In general, animal-borne transmitter devices are designed to achieve high reliability, low weight, small size, and long battery life, giving rise to a complex trade-off.

Among the best transmitters, in terms of battery life, size, and weight, are analog frequency-modulated (FM) radio transmitters. Their low power requirement minimizes the frequency of animal handling and associated handling stress, making them an excellent choice for longitudinal observations of small vertebrates (Ter Maat et al. 2014; Gill et al. 2016; 2015).

Among the challenges associated with FM radio reception is radio signal fading due to relative movements of animal-borne transmitters and stationary receivers. Fading arises when electromagnetic waves arrive over multiple paths and interfere destructively (channel fading) (Tse and Viswanath 2005), e.g., by reflection of metallic walls. Fading also occurs because every receiver has a direction of zero gain, which may affect the reception of the signal from a moving transmitter. Signal fading can be addressed with antenna diversity, i.e., the use of several antennas. To combine the diverse antenna signals, either the strongest signal is selected, all signals are summed up, or signals are first weighted by their strength and then summed (Shatara 2003). However, these approaches do not completely prevent fading because the signals can still annihilate. Alternatively, diversity combining is possible with phase compensation, which is the technique of turning signal phases such that phase compensated signals align and sum constructively (Voitsun, Senega, and Lindenmeier 2020; Senega, Nassar, and Lindenmeier 2017). Phase compensation reduces fading and increases the signal-to-noise ratio of the received signal, and, furthermore, it provides cues for localizing a transmitter (Haniz et al. 2017; Berdanier and Wu 2013). We set out to bring the benefits of antenna diversity and phase compensation techniques to ethology research.

In this report, we present a custom system (BirdPark) that combines all the advantages of the beforementioned multimodal approaches. The purpose of BirdPark is to perform individual-level longitudinal observations of social behaviors. We built a naturalistic environment inside a soundproof enclosure that features a set of microphones to record sounds and several video cameras to capture the entire scene. Moreover, all animals wear a miniature low-power transmitter device that transmits body vibrations from a firmly attached accelerometer via an analog, frequency-modulated (FM) radio signal that we receive with several antennas. The combination of multiple antennas minimizes signal losses and improves the signal-to-noise ratio.

Our system is optimized for robust longitudinal recordings of vocal interactions in songbirds. The on-animal accelerometers enable week-long monitoring of vocalizations without a change of battery. All sensor signals are perfectly synchronized, which we achieve by routing dedicated sample triggers to all recording devices (radio receiver, stationary microphone digitizer, and cameras) derived from a central quartz clock using clock dividers. We release our custom recording software and we demonstrate the high data quality and redundant signaling of vocal gestures.

## Materials & Methods

### The chamber

All our experiments were performed in a sound-isolation chamber (Industrial Acoustic Company, UK) of dimensions 124 cm (width) x 90 cm (depth) x 130 cm (height). The isolation chamber has a special silent ventilation system for air exchange at a rate of roughly 500 l/min. Two types of light sources made of light emitting diodes (LEDs, Waveform Lighting) are mounted on aluminum plates on the ceiling of the chamber: 1) LEDs with natural frequency spectrum (consuming a total electric power of 80 W); and 2) ultraviolet (UV) LEDs of wavelength 365 nm and consuming a total electric power of 13 W.

The ceiling of the chamber contains three circular holes of 7 cm diameter through which we inserted aluminum tubes of 5 mm wall thickness that serve as holders for three cameras (Basler acA2040-120uc). The tubes also conduct the heat of the LEDs (93 W) and the cameras (13 W) to the outside of the chamber where we placed silent fans to keep the cameras below 55 ° C.

With the cameras, we filmed the arena directly from the top and indirectly from two sides via two tilted mirrors in front of the glass side panels. This setup yields a large object distance of about 1 – 1.5 m, which allows for a small perspective distortion.

We installed four microphones on the four side walls of the isolation chamber and one microphone on the ceiling. We mounted two more microphones in the nest boxes and one microphone outside the isolation chamber. Where possible, we covered the sidewalls of the chamber with sound-absorbing foam panels. A door sensor measures whether the door of the chamber is open or closed. The temperature inside the chamber was roughly 26 °C and the humidity was roughly 24%.

The isolation chamber attenuates sounds by roughly 30 dB, which we measured by playing white noise on a loudspeaker inside the chamber and by recording it both inside and outside the chamber. We defined the sound attenuation as the difference in log-RMS amplitude in the 400 Hz to 5 kHz frequency band. The background acoustic noise level inside the chamber is roughly 37 dBA (measured with a Voltcraft SL-10 sound-level meter).

The acoustic reverberation time of the chamber inside is roughly 36 ms. We estimated the reverberation time in terms of the impulse response function that we measured by playing and recording a white noise signal on a loudspeaker inside the chamber. The reverberation time we defined as the linear decay time of the logarithm of the smoothed squared impulse response function over a decay range of 60 dB.

Inside the chamber, we placed the bird arena of dimensions 90 cm x 55 cm (floor). To minimize acoustic resonances and optical reflections, the 40 cm high side panels of the arena are tilted by 11° and 13° toward the inside. Two of the side panels are made of glass for videography and the two opposite panels are made of perforated plastic plates. The floor of the arena is covered with a sound-absorbing carpet. A pyramidal tent made of a fine plastic net covers the upper part of the arena, reaching up to 125 cm above ground. At a height of 35 cm, we attached two nest boxes to the side panels, each equipped with a microphone and a camera. Furthermore, the arena is equipped with perches, a sand-bath, and food and water trays.

### Animals and Experiments

We bred and raised 20 zebra finches (Taeniopygia Guttata): 8 females aged 13-576 dph (day post hatch) and 12 males aged 13-1326 dph in our colony (University of Zurich and ETH Zurich).

The animals were recorded in 6 groups: four 2-bird groups (recording duration 2-15 days), one 4- birds group (recording duration 67 days), and one 8-birds group (recording duration: 89 days).

On each bird, we mounted a transmitter device (Figure 2), and we kept them in the BirdPark on a 14/10 h light/dark daily cycle, with food and water provided ad libitum.

In this study, we were interested in evaluating the feasibility of our behavioral recording methods and in benchmarking their performance, whence the small sizes of the groups with four and eight birds. The experiments were designed to study vocal communication during reproductive behaviors (groups of two birds) and during song ontogeny (groups of four and eight birds). These scientific aims will be pursued in other publications.

All experimental procedures were approved by the Cantonal Veterinary Office of the Canton of Zurich, Switzerland (license numbers ZH045/2017 and ZH054/2020). All methods were carried out in accordance with the Swiss guidelines and regulations concerning animal experiments (Swiss Animal Welfare Act and Ordinance, TSchG, TSchV, TVV). The reporting in the manuscript follows the recommendations in the ARRIVE guidelines.

### Video acquisition system

To visualize the arena, we used industrial cameras (3 Megapixel, MP) with zoom lenses (opening angles: top view: 45 ° x 26 °, back view: 55 ° x 26 °, side view: 26 ° x 35 °) and exposure times of 3 ms. To visualize nests in the dimly lit nest boxes, we used monochrome infrared cameras (2 MP, Basler daA1600-60um) and fisheye lenses (143 ° x 112 °).

The uncompressed camera outputs (approx. 400 MB/s in total) are relayed on a host computer via a USB3 Vision interface. Each camera receives frame trigger signals generated by the USRP ((Chen, Matheson, and Sakata 2016; Carouso-Peck and Goldstein 2019), C). A custom program (BirdVideo), written in C++ and using the OpenCV library (Gary Bradski 2000) and FFMPEG (Tomar 2006), undistorts the nest camera images with a fisheye lens model and transforms the five camera images into a single 2976 x 2016 pixel-sized image (Figure 1B). The composite images are then encoded with the h264 codec on a NVIDIA GPU and stored in an MP4 (ISO/IEC 14496-14) file. We used constant-quality compression with a variable compression ratio in the range 150-370, resulting in a data rate of 0.72 MB/s – 1.8 MB/s, depending on the number of birds and their activity in the arena. Compression ratios in this range do not significantly decrease key point tracking performance (Mathis and Warren 2018). The frame rate of the video is about 48 frames per second (frame period 21 ms). The spatial resolution of the main cameras is about 2.2 pixels/mm.

**Figure 1:**
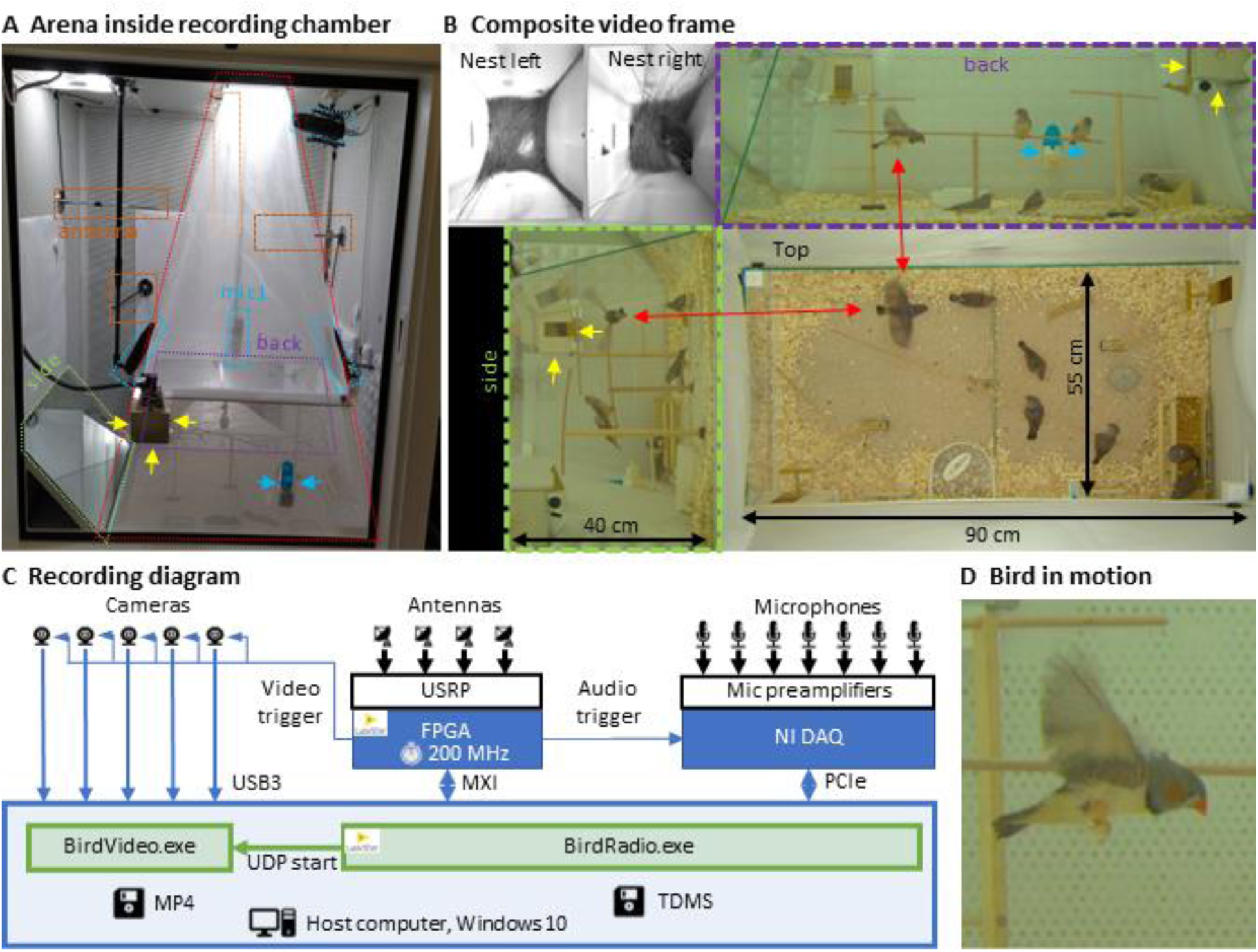
Recording arena and recording system schematic. (**A**) Inside a soundproof chamber, we built a recording arena (red dotted line) for up to 8 birds. We record the animals’ behaviors with three cameras mounted through the ceiling. These provide a direct top view and indirect side and back views via two mirrors (delimited by green and purple dotted lines). To record the sounds in the chamber, we installed five microphones (blue dotted lines) among all four sides of the cage (one attached to the front door is not visible) and the ceiling, and two small microphones in the nest boxes. The radio signals from the transmitter devices are received with four radio antennas (orange dotted lines) mounted on three side walls and the ceiling. One nest box is indicated with yellow arrows and a water bottle with blue arrows. (**B**) A composite still image of all camera views shows two monochrome nest box views (top left) and three views of the arena (top, side, back) with 8 birds among which one is flying (red arrows). Yellow and blue arrows as in A. (**C**) Schematic of the recording system for gapless and synchronized recording of sound (microphones), acceleration (transmitter devices), and video channels (cameras). The radio receiver is implemented on a universal software radio peripheral (USRP) with a large field programmable gate array (FPGA) that runs at the main clock frequency of 200 MHz. Clock dividers on the FPGA provide the sample trigger for audio recordings and the frame trigger for the cameras. The data streams are collected on a host computer that runs two custom programs, one (BirdRadio) for streaming audio and sensor signals to disk and one (BirdVideo) for encoding video data. (**D**) Zoom-in on an airborne bird, illustrating the spatial and temporal resolution of the camera.

**Figure 2:**
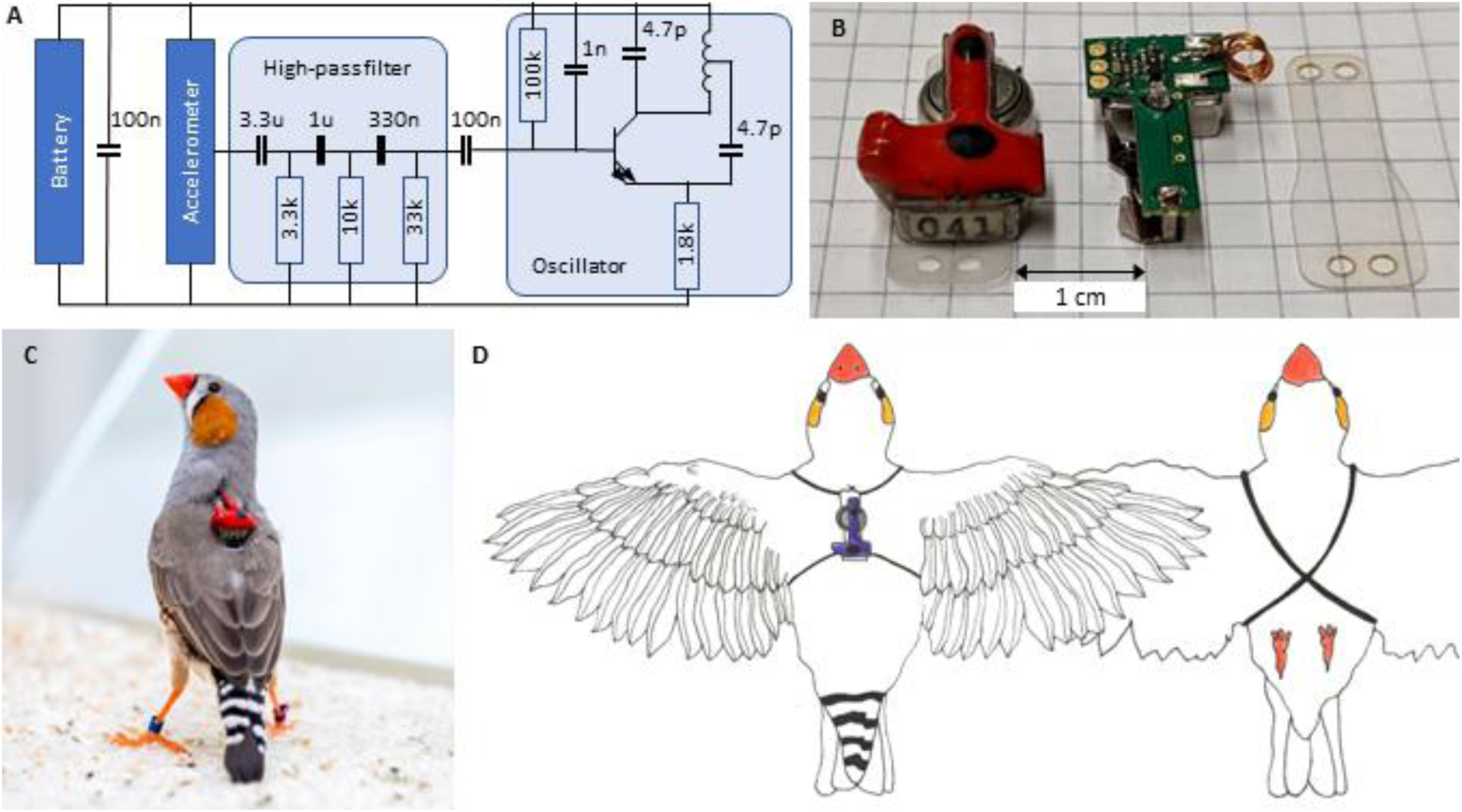
Transmitter device. (A) Schematic of the electronic circuit (adapted from Ter Maat et al. (2014)). The analog FM radio transmits the vibration transducer (accelerometer) signal via a high-pass filter (cutoff frequency: 15Hz) followed by a radiating oscillator. (B) A fully assembled transmitter (left), and another one without epoxy and battery (middle), and a piece of mounting foil (right). (C) Picture of the device mounted on a bird. The transmitters are color-coded to help identify the birds in the video data. (D) Schematic of a bird wearing the device. The harness (adapted from Alarcón-Nieto et al. (2018)) is made of rubber string (black) that goes around the wings and the chest.

### Transmitter device

The accelerometer (Knowles BU21771) senses acceleration with a sensitivity of 1.4 mV/(m/s2) in the 30 Hz – 6 kHz frequency range (Knowles Electronics 2017). Inspired by Ter Maat et al. (2014) we performed the frequency modulation of the accelerometer signal onto a radio carrier using a simple Hartley oscillator transmitter stage with only one transistor and a coil that functions both as the inductor in the LC resonator and as the antenna. The LC resonator determines the carrier frequency 𝜔_*c*_ of the radio signal. We set 𝜔_*c*_ of our transmitters in the vicinity of 300 MHz, which corresponds to an electromagnetic wavelength of about 1 m and is roughly equal to the dimension of our soundproof chambers, implying that our radio system operates in the near field.

We found that 𝜔_*c*_ depends on temperature at an average of -73 kHz/°C (range -67 kHz/°C to -87 kHz/°C, n=3 transmitters). Towards the end of the battery life (over the course of the last 3 days), we observed an increase of 𝜔_*c*_by an average of 500 – 800 kHz. These slow drifts can easily be accounted for by tracking and high-pass filtering the momentary frequency. The measured end-to-end sensitivity of the frequency modulation is 5 kHz/g, with g being the gravitational acceleration constant.

The transmitters are powered by a zinc-air battery (type LR41), which lasts at least 12 days. The total weight of the transmitter including the harness and battery is 1.5 g, which is ca. 10% of the body weight of a zebra finch.

We moderately tightened the harness during attachment in a tradeoff between picking up vibratory signals from the singer and preserving the wearer’s natural behaviors.

### Radio reception

We mounted four whip antennas of 30 cm length (Albrecht 6157) perpendicular to the metallic sidewalls and ceiling of the chamber (using magnetic feet). We fed the antenna signals to a universal software radio interface (USRP-2945, National Instruments, USA), which comprises four independent antenna amplifiers the gains of which we set to 68 dB, adjustable in the range - 20 dB to +90 dB.

### Intermediate band

The USRP generates from the amplified radio signal a down-converted signal with an analog bandwidth of 80 MHz around the local oscillator frequency (ω_*LO*_). After digitization, this intermediate signal *Z*^a^(*t*) for antenna a ∈ {A, B, C, D}, is a 100 MS/s complex-valued signal (IQ) of 2 x 18 bits precision and is relayed to the FPGA. For details about the analog down conversion and the sampling of complex valued signals, see the NI manual of the USRP and the documentation of our custom radio software.

### Digital down-converters (DDCs) and baseband

The PLL of channel _i_ ∈ {1, . . .,8} makes use of four digital down-converters (DDCs, Figure 3D), one for each antenna a, that cut from the intermediate band a narrow baseband around the tracking frequency ω_*i*_. The 3-stage decimation filter of this down-conversion has a flat frequency response within a ±100 kHz band. Given the decimation factor of 128, the resulting complex base-band signals 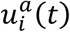 (the outputs of the DDCs) have a sample rate of 781.25 kHz and a precision of 2 x 25 bits.

**Figure 3:**
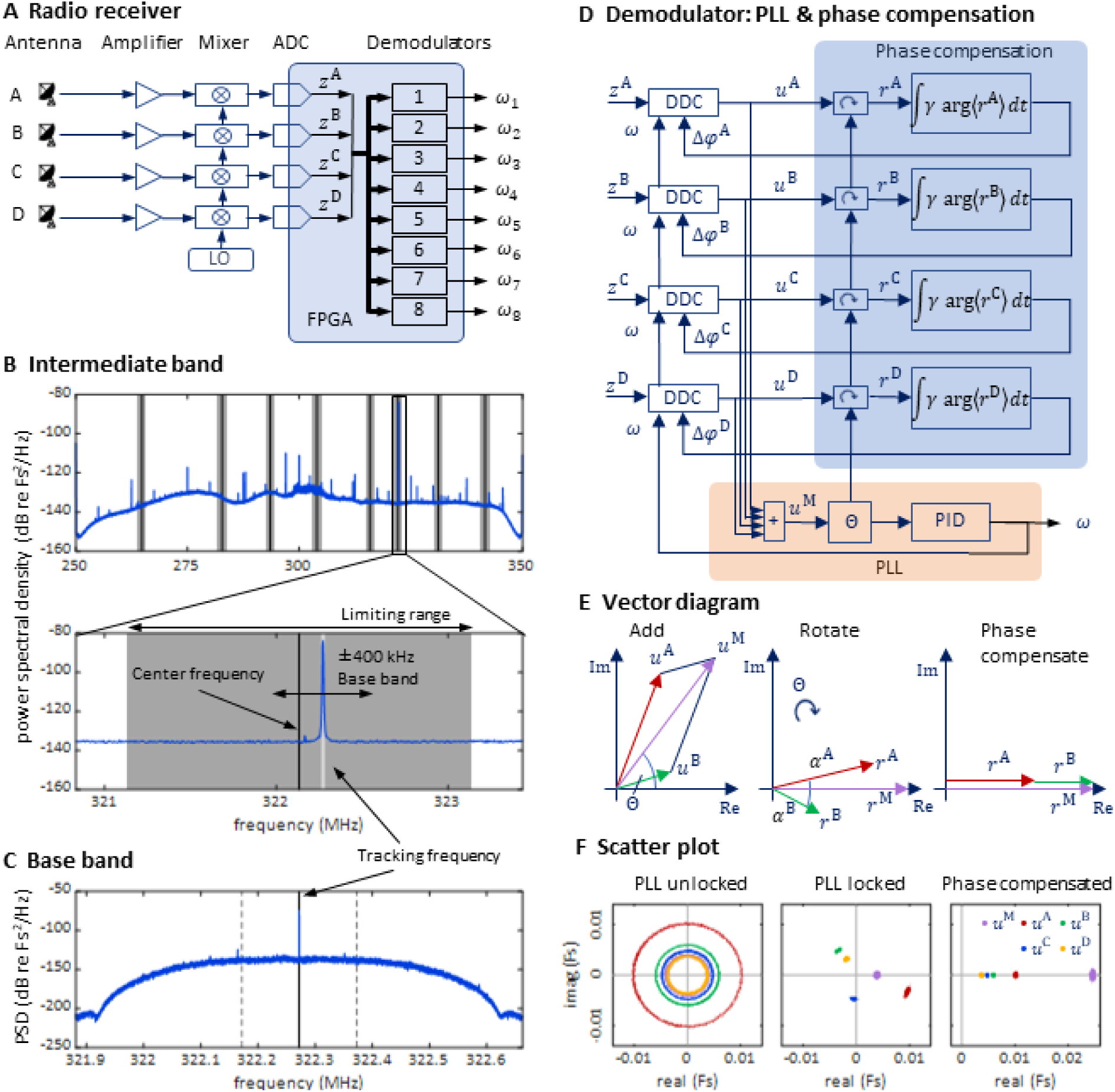
Radio receiver with PLL demodulators and phase compensation. (**A**) Schematic of the software-defined radio receiver. Analog antenna signals are first down converted by mixing the amplified signals with a local oscillator of frequency 𝜔_*LO*_. The resulting intermediate signals have digital representations *Z*^*A*^(*t*), *Z*^*B*^(*t*), *Z*^*C*^(*t*), and *Z*^*D*^(*t*). From these, eight FM demodulators extract the transmitter tracking frequencies 𝜔_*i*_(*t*), *i* ∈ {1, . . .,8}. (**B**) The power spectral density of the 80 MHz wide intermediate band *Z*^*A*^(𝜔) centered on the oscillator frequency 𝜔_LO_ =300 MHz. The limiting ranges (gray bars) of the 8 channels (transmitters) are typically set to ± 1 MHz of their manually set center frequencies (black vertical lines). A zoom-in (lower graph) to the limiting range of one channel reveals the tracking frequency 𝜔(*t*) (light grey vertical line) that tracks the large peak at the transmitter frequency. (**C**) The baseband power spectral density *u*^*A*^(𝜔) is a down-converted, ±100 kHz wide (dashed vertical lines), flat band around the tracking frequency (solid vertical line) close to the transmitter frequency (the large peak). (**D**) Schematic of one demodulator: a PLL (orange shading) computes the tracking frequency 𝜔 and four phase compensation circuits (blue shading) align the baseband signals *u*^a^(*t*) which are derived from intermediate signals with digital down-converters (DDCs) operating on a common tracking frequency 𝜔. (**E**) Vector diagram in the complex plane illustrating the effects of the PLL (middle) and of phase compensation (right) on main and baseband vectors (shown only antenna A & B). The phase compensation circuits drive the phases 𝛼^a^ = a𝑟𝑔(⟨𝑟^a^⟩) of rotated baseband signals 𝑟^a^ = *u*^a^𝑒^−*i*𝜃^ to zero. (**F**) The four baseband vectors *u*^a^ are shown without phase compensation (left), after aligning the main vector with the PLL (middle), and after additional phase compensation (right). The result is that all vectors are aligned, and the master vector is of maximal amplitude.

### Phase locked loop

Each PLL generates its tracking frequency ω_*i*_ by direct digital synthesis (DDS) with a 48-bit phase accumulation register and a lookup table. This frequency is dynamically adjusted to keep the phase of the main vector (i.e., the summed baseband signals) close to zero. We calculated the angle 𝜃_*i*_ = arg(*u*^M^) of the main vector with the CORDIC algorithm (Volder 1959) and unwrapped the phase up to ±128 turns before using it as the error signal for a proportional- integral-derivative (PID) controller. The PID controller adjusts the tracking frequency of the PLL.

The PID controller was implemented on the FPGA in single precision floating point (32-bit) arithmetic. The controller included a limiting range and an anti-windup mechanism (Astrom and Rundqwist 1989). The unwrapping of the error phase was crucial for the PLL to quickly lock-in and to keep the lock even during large and fast frequency deviations of the transmitter. To tune the PID parameters, we measured the closed-loop transfer function of the PLL by adding a white-noise signal to the control signal (tracking frequency), and then we adjusted the PID parameters until we observed a closed loop transfer function with low-pass characteristic. We achieved a closed loop bandwidth of about 30 kHz.

### Phase compensation

For a given PLL *i*, we compensated the relative phases under which a radio signal arrives at the four antennas a ∈ {A, B, C, D} in order to align the four baseband vectors 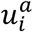. The alignment was achieved by providing a phase offset 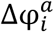 to each down-converter, where it acts as offset to the phase accumulation register of the direct digital synthesis. To compute the angle of baseband vector relative to the main vector, we rotated the baseband vectors 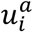 by the phase 𝜃_*i*_ of the main vector to result in the rotated vector 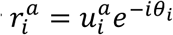. After averaging the rotated vectors across 512 samples, we computed its angle 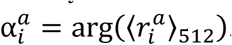. We then compensated that angle by iteratively adding a fraction γ ∈ [0,1] of it to the phase offset: 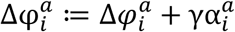. The parameter γ is the phase compensation gain (Figure 3D), typically set to 𝛾 = 0.2.

Because birds’ locomotion is slower than their rapid vocalization-induced vibratory signals, the phases 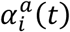 change more slowly than the tracking frequency 𝜔_*i*_(*t*) and therefore we updated phase offsets less often than the PLL’s tracking frequency at a rate of about 1.5 kHz (781.25 kHz / 512).

### Central control software

On the host computer we run our central control software (BirdRadio) programmed with LabView that acquires the microphone and transmitter signals and writes them to a TDMS file (Figure 1C). Furthermore, BirdRadio sends UDP control signals to BirdVideo, it automatically starts the recording in the morning and stops it in the evening, it controls the light dimming in the sound-isolation chamber with simulation of sunrise and sunset, it triggers an email alarm when the radio signal from a transmitter device is lost, and it automatically adjusts the center frequency of each radio channel every morning to adjust for carrier frequency drift.

### Data management

The BirdPark is designed for continuous recordings over multiple months, producing data at a rate of 60 GB/day for 2 birds and 130 GB/day for 8 birds. We implemented the FAIR (findable, accessible, interoperable, and reusable) principles of scientific data management (Wilkinson et al. 2016) as follows: Our recording software splits the data into gapless files of 20’000 video frames (ca. 7 mins duration). At the end of a recording day, all files are processed by a data compilation script that converts the TDMS files into HDF5 files and augments them with rich metadata. The HDF5 files are self-descriptive in that they contain metadata as attributes and additionally, every dataset in the file contains its description as an attribute. We use the lossless compression feature of HDF5 to obtain a compression ratio of typically 2.5 for the audio and accelerometer data. The script also adds two AAC-compressed microphone channels into the video files. Although this step introduces redundancy, the availability of sound in the video files is very useful during manual annotation of the videos. Furthermore, the script also exports the metadata as a JSON file and copies the processed data onto a NAS server. At the end of an experiment, the metadata is uploaded onto an openBIS (Bauch et al. 2011) server and is linked with the datafiles on the NAS.

### Radio signal-to-noise ratio (RSNR)

We calculated the 𝑅𝑆𝑁𝑅^a^(*t*_𝑘_), a ∈ {A, B, C, D, M} of the non-overlapping 𝑘-th radio frame at center time *t*_𝑘_ as:

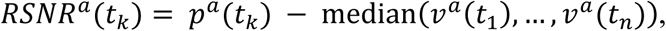

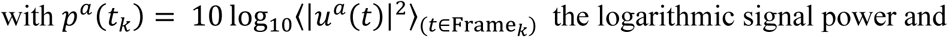

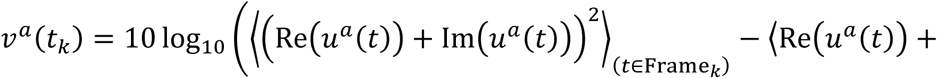

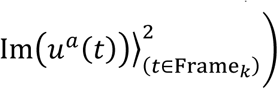 the logarithmic noise power, where the averages are calculated over radio frames and the median is calculated over 7-mins long files (n=20 000 radio frames).

We compared the multi-antenna 𝑅𝑆𝑁𝑅^M^(*t*_𝑘_) to the 𝑅𝑆𝑁𝑅^∗^(*t*_𝑘_) of the best single-antenna ∗defined as:

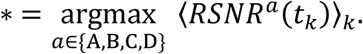

### Manual segmentation of vocalizations

We manually segmented vocalizations in a randomly chosen file of 7 mins duration of two experiment instances with two birds (copExpBP08 and copExpBP09) and in the first 1 min of seven randomly chosen files in an experiment instance with four birds (juvExpBP01) and one with eight birds (juvExpBP03). We excluded files with a signal loss rate above 0.1%. We high-pass filtered (a passband frequency of 200 Hz) the raw demodulated transmitter signals and microphone signals and produced spectrograms (window size=384 samples, hop size=96 samples) that we manually segmented using Raven Pro 1.662 or BpBrowser (our custom data annotator). We performed three types of segmentations in the following order:

1. Transmitter-based vocal segments: On all transmitter channels, we separately segmented all vocalizations, precisely annotating their onsets and offsets. In the released data sets, these segments are referred to as transmitter-based vocal segments. We visually checked for crosstalk (whether we saw a faint trace of a vocalization on another bird’s transmitter spectrogram). We excluded crosstalk segments from further analysis (except for the crosstalk statistics): If a segment was detected as crosstalk, we set the Tag Crosstalk to the transmitter channel name of the bird that generated the crosstalk, otherwise we set the tag to ‘No’. When we were uncertain whether a sound segment was a vocalization or not, we also looked at spectrograms of microphone channels and listened to sound playbacks. If we remained uncertain, we tagged the segment with the label ‘Unsure’: we treated such segments as (non-vocal) noises and excluded them from further analysis (except for the uncertainty statistics).
2. Microphone-based vocal segments: We simultaneously visualized all microphone spectrograms (we ignored nest mics when the nest was not accessible to the birds) using the multi-channel view in Raven Pro (we ignored Mic7, which was located in the second nest that was not accessible to the birds). On those, we annotated each vocal segment on the first microphone channel on which it was visible (e.g., a syllable that is visible on all microphone channels is only annotated on Mic1). Overlapping vocalizations were annotated as a single vocal segment. When we were uncertain whether a sound segment was a vocalization or not, we also looked at spectrograms of transmitter channels and listened to sound playbacks. If we remained uncertain, the segment was tagged with the label ‘Unsure’, such segments were treated as (non-vocal) noises and were excluded from further analysis.
3. Consolidated vocal segments: All consistent (perfectly overlapping down to a temporal resolution of one spectrogram bin) transmitter- and microphone-based vocal segments, we labelled as consolidated vocal segments. We then inspected all inconsistent (not perfectly overlapping) segments by visualizing all channel spectrograms. We fixed inconsistencies that were caused by human annotation errors (e.g. lack of attention) by fixing the erroneous or missing transmitter- and microphone-based segments. From the inconsistent (partially overlapping) segments that were not caused by human error, we generated one or several consolidated segments by trusting the modality that more clearly revealed the presence of a vocalization (hence our reporting of ‘misses’ in Supplementary Table S1).

In our released annotation table, we give each consolidated vocal segment a Bird Tag (e.g., either ‘b15p5_m’ or ‘b14p4_f’) that identifies the bird that produced the vocalization, a Transmitter Tag that identifies the transmitter channel on which the vocalization was identified (either ‘b15p5_m’ or ‘b14p4_f’ or ‘None’), and a FirstMic Tag that identifies the first microphone channel on which the segment was visible (‘Mic1’ to ‘Mic6’, or ‘None’). We resolved inconsistencies and chose these tags as follows:

- If a microphone (-based vocal) segment was paired (partially overlapping) with exactly one transmitter segment, a consolidated segment was generated with the onset time set to the minimum onset time and the offset time set to the maximum offset time of the segment pair. The Bird and Transmitter Tags were set to the transmitter channel name, and the FirstMic Tag was set to the microphone channel name.
- If a transmitter segment was unpaired, a consolidated segment was created with the same onset and offset times. The Bird and TrCh Tags were set to the transmitter channel name, and the FirstMic Tag was set to ‘None’.
- If a microphone segment was unpaired, we tried to guess a transmitter channel name based on the vocal repertoire and noise levels on both transmitter channels. We visually verified that the vocal segment was not the result of multiple overlapping vocalizations (which was never the case). Then we created a consolidated segment with the same on- and offset, set the FirstMic Tag to the microphone Id, the Bird Tag to the guessed transmitter channel name or ‘None’ if we were unable to make a guess, and the Transmitter Tag to ‘None’.
- If a microphone segment was paired with more than one transmitter segment, a consolidated segment was created for each of the transmitter segments. The onsets and offsets were manually set based on visual inspection of all spectrograms. Bird and Transmitter Tags were set to the transmitter channel name and the FirstMic Tag was set to the first microphone channel on which the respective segment was visible.
- We never encountered the case where a transmitter segment was paired with multiple microphone segments.

The statistics in Supplementary Table S1 were calculated as follows: # vocalizations = (number of consolidated segments)

\# voc. missed on tr. channel = (number of consolidated segments with Transmitter=None)

\# voc. missed on all mic. channels = (number of consolidated segments with FirstMic=None)

\# voc. missed on Mic1 = (number of consolidated segments with FirstMic≠Mic1 and FirstMic≠None)

\# voc. unassigned = (number of consolidated segments with Bird=None)

\# voc. with overlaps = (number of consolidated segments which overlap with at least one other syllable by at least one spectrogram bin)

\# voc. with crosstalk = (number of transmitter-based segments with Crosstalk)

\# uncertain segments = (number of transmitter-based and microphone-based segments with ‘Unsure’ tag)

### Radio frequency jumps

We detected frequency jumps as deviations of the transmitter tracking frequency ω_i_larger than 50 kHz from the running median computed within 0.2 s sliding windows. Frequency jumps were concatenated when they were separated by less than 50 ms. The modulation amplitude was defined as the maximum absolute deviation.

### Wing Flap Event Detection and Video Alignment

The transmitter signals evoked by wing flap movement are downward and dip-like (Figure 4). We detected dips on the male’s transmitter signal by thresholding its derivative after down- sampling by a factor of 32. A wing flap event is recognized when the derivative initially falls below a negative threshold, −𝑊 = −60 MHz / s, and subsequently rises above a positive threshold, 𝑊 within a 2 to 12 ms window following the initial threshold crossing. The detected dips correspond closely with the center time point of the wing’s down stroke (Figure 7B).

**Figure 4:**
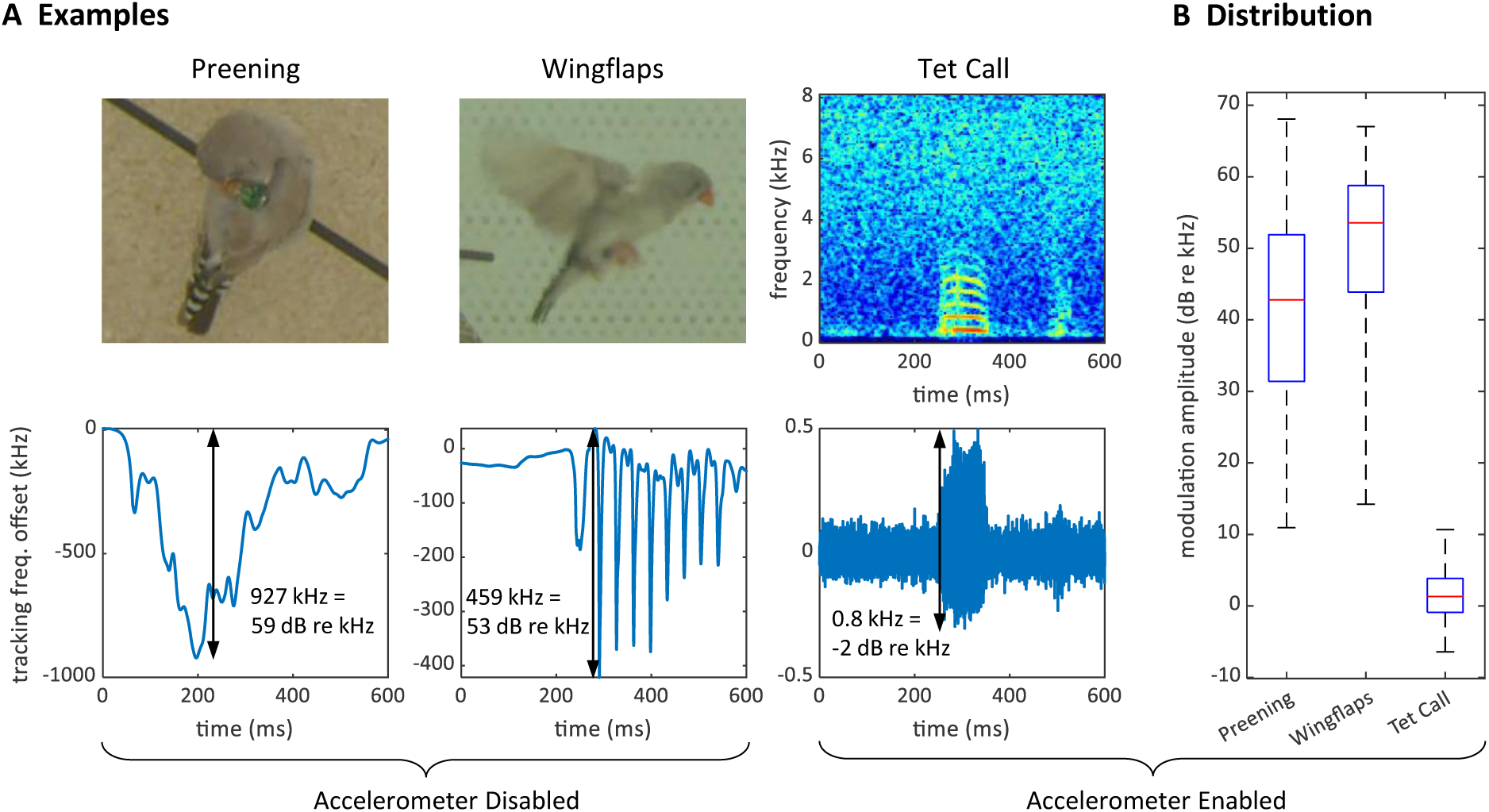
Large proximity effects on the transmitter frequency. (**A**) On a transmitter device with a disabled (short circuited) accelerometer, the transmitter tracking frequency is strongly modulated by the proximity of the head during preening (left) and by wing movements during flight (middle). In contrast, the modulation of the (high-pass filtered) tracking frequency is much weaker for vocalizations (right) on a transmitter with enabled accelerometer. The modulation amplitudes (black arrows) due to proximity are about 1000 times larger than the vocalization-induced modulation amplitude. (**B**) Distribution of modulation amplitudes for preening (n=62), wing flaps (n=54), and tet calls (n=105). The preening and wing flap events were manually annotated from a set of randomly selected concatenated frequency jumps defined as data segments associated with absolute tracking frequency jumps larger than 50 kHz (see methods) and taken from a 25-h long recording of a bird with a transmitter with a disabled accelerometer. Tet call segments that were not masked by noise were manually annotated on datafiles of a large dataset of mixed-sex zebra finch pairs.

We time-stamped the video frames as follows. For each detected wing-flap event, we examined the transmitter’s signal range. Events associated with a signal range less than 200 kHz we discarded from further analysis, where we defined the signal range as the maximum minus the minimum transmitter signal in a time window [-10.5, 10.5] ms (corresponding to 1 video frame period) centered on the minimum of the signal dip. Similarly, we excluded abrupt signal changes exceeding 1000 kHz.

The video frame with exposure onset nearest to the dip minimum we defined as the first event frame, and the subsequent frame as the second event frame. We then ordered all event frames according to their time stamp: the time lag between the exposure onset and the dip minimum.

By randomly sampling video frames in which the flapping birds were clearly visible and continuously flying (from n=5 birds), we obtained the illustration of wing flapping shown in Figure 7, nicely capturing the phases of the wing flap cycle. All the frames in Figure 7 are derived from a comprehensive dataset of 5 birds, with each event capturing a wing flap from a randomly selected bird.

## Results

### The Recording Arena

We built an arena optimized for audiovisual recordings, minimizing acoustic resonances and visual occlusions (Figure 1A). It provides space for up to 8 songbirds and contains nest boxes, perches, sand baths, food, and water sources. To record the sounds inside the chamber, we installed five microphones. Three video cameras capture the overall scene from three orthogonal viewpoints. In addition, we installed a camera and microphone in each of the two nest boxes. To simplify video analysis, we combined the images from all five cameras into a single video image (Figure 1B). Camera resolution is high enough and exposure times short enough to resolve key points on birds even in mid-flight (Figure 1D).

To record vocalizations, we mounted transmitter devices to birds’ backs. Each device comprises an accelerometer that picks up body vibrations from vocalizations and body movements such as hopping and wing flapping (Anisimov et al. 2014). The devices transmit the accelerometer signals as frequency-modulated (FM) radio waves to four antennas inside the recording chamber.

We demodulated the FM radio signals with a custom eight-channel radio receiver that we implemented on a universal software radio peripheral (USRP) with a large field programmable gate array (FPGA). To ensure that the data streams from the video cameras, microphones, and transmitter devices are synchronized, we record videos with industrial cameras and digitize sounds on a National Instruments data acquisition board (NI DAQ), both of which receive sample trigger signals from the USRP. The base clock of the USRP was 200 MHz and was divided by 213 to generate the audio sample rate of 24.414 kHz, and with a further division by 29, we generated the 47.684 Hz video frame rate.

On the host computer that controlled the acquisition system, we ran two custom applications: BirdVideo, which acquires and writes the video data to a file, and BirdRadio, which acquires and writes the microphone and transmitter signals to a file (see Methods). The generated file pairs are synchronized such that for each video frame there are 512 audio samples (an audio frame) and 512 transmitter signal samples (a radio frame). While the recording is gapless, it is split into files of typically 7 minutes duration.

### Transmitter device

Our transmitter devices are based on the FM radio transmitter circuit described in Ter Maat et al. (Figure 2A). To distinctly record birds’ vocalizations irrespective of external sounds, we replaced the microphone with an accelerometer (Anisimov et al. 2014) that picks up body vibrations and acts as a contact microphone. The radio circuit uses a single transistor to modulate the sensor signal a_𝑇_(*t*) (where 𝑇 stands for Transmitter and *t* is the time) onto a radio carrier frequency 𝜔_*c*_(*t*) (set by the resonator circuit properties) and has an inductor coil as an emitting antenna. As a result, the accelerometer signal a_𝑇_(*t*) is encoded as (instantaneous) transmitter frequency 𝜔_𝑇_(*t*) ≃ 𝜔_*c*_(*t*) + *c*a_𝑇_(*t*) with *c* some constant.

Before mounting a device on a bird, we adjusted the coil to the desired carrier frequency in the range of 250–350 MHz by slightly bending the coil wires. Thereafter, we fixed the coil and electronics in dyed epoxy. The purpose of the dyed epoxy is to help identify the birds in the video images, protect the printed circuit board, and stabilize the coil’s inductance (Figure 2B, C). We mounted the devices on birds using a rubber-string harness adapted from Alarcón-Nieto et al.(2018) (Figure 2D).

### Radio receiver

To reconstruct the accelerometer signal a_𝑇_(*t*) of a given transmitter device from the received multi-antenna FM signals, we demodulated the latter using a phase-locked loop (PLL), which measures the instantaneous transmitter frequency 𝜔_𝑇_(*t*). A PLL generates an internal oscillatory signal of variable tracking frequency 𝜔(*t*). It adjusts that frequency to maintain a zero phase with respect to the received (input) signal. As long as the zero-phase condition is fulfilled, the PLL’s tracking frequency 𝜔(*t*) follows the instantaneous transmitter frequency 𝜔_𝑇_(*t*). The high-pass filtered tracking frequency 𝜔_𝛼_(*t*) is our estimate of the accelerometer signal a_𝑇_(*t*). We refer to it as the (demodulated) transmitter signal.

In our diversity combining approach, we construct the PLL’s input signal as a combination of all four antenna signals. Namely, we compensate the individual phase offsets of the four antenna signals in such a way that all phases are aligned (we achieve this phase shifting with phase-compensation circuits — one for each antenna). We then form the desired mixture signal by summing the phase-shifted signals; the phase of the summed signal serves as the PLL’s error signal that we use to adjust the tracking frequency 𝜔(*t*). The variable phase offsets on the four antennas arise from the variable locations and orientations of transmitters relative to receivers.

In summary, our approach to minimizing fading in wireless ethology research is to use a demodulator comprising a PLL and a diversity combining approach based on phase-compensation. Our FM radio demodulation is described in more detail in the following.

We implemented our custom demodulation technique by installing four antennas labeled A, B, C, and D perpendicular to the walls and the ceiling of the chamber (see Methods: Radio reception). We fed the antenna signals into a USRP containing a large FPGA (Figure 3A). The four input stages of the USRP filter out an 80 MHz wide band around the local oscillator frequency ω_LO_, which we typically set to 300 MHz. These four signals then become digitally available on the FPGA as complex valued signal of 100 MHz sampling rate. We call them the intermediate signals *Z*^a^(*t*), a ∈ {A, B, C, D} (intermediate band in the frequency domain, Methods: Intermediate band, Figure 3B).

On the FPGA, we instantiated eight demodulators that simultaneously demodulated up to eight transmitter channels. Each demodulator contained four digital down-converters (DDC) that extracted from the four intermediate signals *Z*^a^(*t*) the four baseband signals *u*^a^(*t*), a ∈

{A, B, C, D} around the tracking frequency 𝜔(*t*) (Figure 3C and D, Methods: Baseband). The digital signals are complex valued, so we interchangeably call them vectors and their phases we refer to as angles.

The PLL is driven by the main vector *u*^M^(*t*) = *u*^A^(*t*) + *u*^B^(*t*) + *u*^C^(*t*) + *u*^D^(*t*) that we formed as the sum of the four baseband vectors. The error signal used in the PLL’s feedback controller is given by the phase of the main vector 𝜃(*t*) = arg(*u*^M^(*t*)) (see Methods: PLL, Figure 3D).

When that phase is approximatively zero (PLL is locked), the tracking frequency 𝜔(*t*) tracks the transmitter frequency.

To avoid destructive interference in the summation of the baseband vectors, we compensated their phases 𝛼^a^ (t) = ∡(*u*^M^ (t), *u*^a^ (t)), a ∈ {A, B, C, D} relative to the main vector. We introduced individual phase offsets Δφ^a^(*t*) that were set in feedback loops to drive the phases 𝛼^a^(*t*) towards zero (Methods: Phase compensation, Figure 3E).

The PLL and phase compensation form independent control loops. When the PLL is unlocked (e.g., off), the tracking frequency does not match the instantaneous transmitter frequency and the baseband vectors rotate at the difference frequency (Figure 3F left). When the PLL is switched on and locked, the baseband vectors do not rotate, and the main signal displays a phase 𝜃(*t*) ≃ 0 (Figure 3F middle). When the phase alignment is switched on, the baseband signals align, and their sum maximizes the magnitude of the main vector (Figure 3E right).

### Operation

The intermediate band of *Z*^a^(𝜔) is wide enough (80 MHz) to accommodate up to eight FM transmitters. Provided the transmitters’ FM carrier frequencies are roughly evenly spaced, the transmitter frequencies do not cross, even during very large frequency excursions. Nevertheless, we limited the tracking frequency ω_*i*_ of channel *i* to the range [Ω_*i*_ − ΔΩ, Ω_*i*_ + ΔΩ], where Ω_i_ is the center frequency and ΔΩ = 1 MHz is the common limiting range of all channels (Figure 3B). The center frequency Ω_*i*_ of a channel we manually set at the beginning of an experiment to the associated FM carrier frequency. The limiting range of 1 MHz we found narrow enough for the PLL to rapidly relock after brief signal losses.

During PLL tracking losses, the tracking frequency ω_*i*_(*t*) almost always stuck to one of the two limits of its range. Therefore, we detected tracking loss events heuristically as the time points *t* satisfying |ω_*i*_(*t*) − Ω_i_ | = ΔΩ. Tracking losses were measured in 14-h time periods per bird group, they occurred at an average rate of 1.07 % (range 0.00 % – 4.50 %, 𝑛 = 5 groups, Table 1).

**Table 1:** Signal loss statistics. Percent time (per radio channel) the signal is lost due to insufficient PLL tracking or signal fading. The latter is reported for both multi-antenna and best single-antenna demodulation. The statistics are reported for the test experiment in Figure 5 (rows 2 and 3) and four additional experiments (copExp and juvExp). The average percentages in the last row exclude the experiment with the complex antenna because of incompatible experimental condition.

| Experiment | # birds | Insufficient PLL tracking (%) | Fading signal: Multi antenna (%) | Fading signal: Best single antenna (%) |
| --- | --- | --- | --- | --- |
| TestExp (complex antenna) | 2 | 0.41 |  | 0.01 |
| TestExp (multi antenna) | 2 | 0.00 | 0.00 | 0.62 |
| copExpBP08 | 2 | 0.67 | 0.00 | 3.44 |
| copExpBP09 | 2 | 0.08 | 0.00 | 2.22 |
| juvExpBP01 | 4 | 0.09 | 0.00 | 4.41 |
| juvExpBP03 | 8 | 4.50 | 0.00 | 2.84 |
| Average (excl. complex ant.) |  | 1.07 | 0.00 | 2.71 |

During operation, we often observed large excursions of the tracking frequency 𝜔(*t*). These excursions occurred while birds pecked on the device or while they preened their feathers near the sensor, or when one bird sat on top of another, such as during copulations. The magnitude of these frequency excursions could reach 2.5 MHz (68 dB re 1 kHz), which is much larger than the maximal 3.4 kHz (11 dB re 1 kHz) shifts induced by vocalizations (proximity effect, Figure 4).

The significant fluctuations in the tracking frequency were likely caused by dielectric effects resulting from body movements near the resonator circuit. Specifically, the proximity of body parts altered the permittivity of the air surrounding the coil, thereby modulating the carrier frequency 𝜔_*c*_. Additionally, the carrier frequency 𝜔_*c*_was subject to temperature and end-of- battery-life drift (see Methods: Transmitter device). While the instability of the carrier frequency 𝜔_*c*_ is a disadvantage of the single-transistor circuit design, advantages are the low device weight (1.5 g) and the small power consumption it enables (the battery lifetime was about 12 days).

### Performance of diversity combining technique

We validated the robustness of our diversity-combining multi-antenna demodulation method by analyzing the signal-to-noise ratio of the received transmitter signal while recording a zebra finch pair over a full day (2 x 14 h measurement period). We quantified the performance by comparing the radio signal-to-noise ratio (𝑅𝑆𝑁𝑅^M^) of the diversity combining multi-antenna signal (M) with the 𝑅𝑆𝑁𝑅^a^ of the four single-antenna signals a ∈ {A, B, C, D}. The 𝑅𝑆𝑁𝑅 was defined as the logarithmic signal power minus the logarithm of the estimated noise variance (see Methods for details). Our recording software BirdRadio logs power and variance trajectories of *u*^a^(*t*), which allows us to evaluate 𝑅𝑆𝑁𝑅^M^and 𝑅𝑆𝑁𝑅^a^ on the same dataset. Furthermore, 𝑅𝑆𝑁𝑅 is a good measure of the signal quality of the demodulated transmitter signal because the noise power of the accelerometer signal 𝜔_𝛼_(*t*) is a monotonically decreasing function of RSNR (Figure 5A, B).

**Figure 5:**
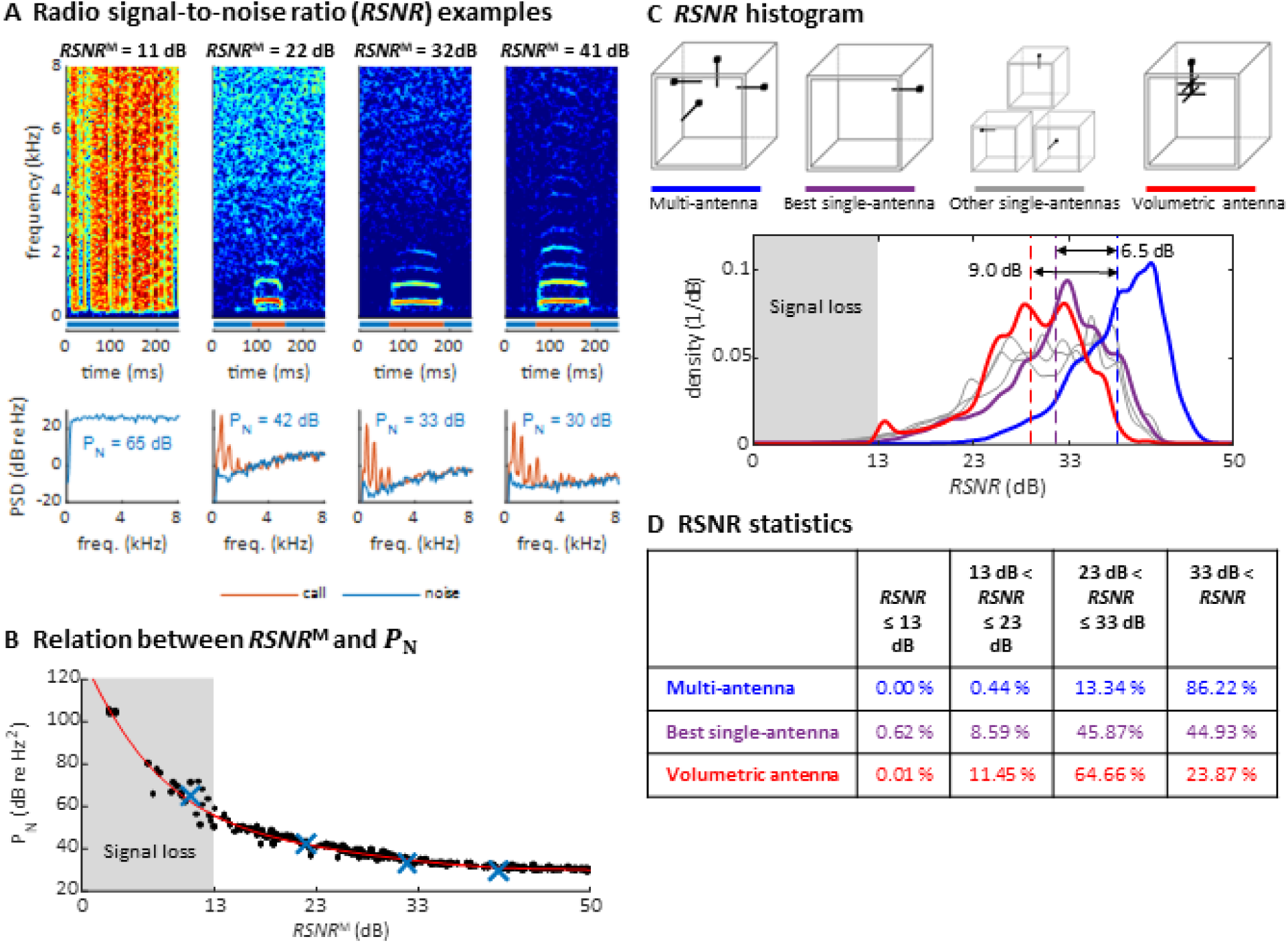
Multi-antenna demodulation achieves the largest radio signal-to-noise ratio (RSNR). (A) Examples of acceleration spectrograms that originate from a single calling zebra finch, with 𝑅𝑆𝑁𝑅^𝑀^s of 11, 22, 32, and 41 dB (top, from left to right). The bottom plots show the spectra of the accelerometer signal during calls (red lines) and noise (blue lines), computed as time averages from the spectrograms above (time segments indicated by red and blue horizontal bars). The noise power 𝑃_𝑁_(integral of blue curve, units: dB re 1 Hz^2^), is decreasing with 𝑅𝑆𝑁𝑅. When the 𝑅𝑆𝑁𝑅 is below 13 dB, the noise power spectral density is above the signal power of most vocalizations (these become invisible). (B) Scatter plot of 𝑃_𝑁_ versus 𝑅𝑆𝑁𝑅^𝑀^ measurements (dots), plotted for noise segments of diverse 𝑅𝑆𝑁𝑅^𝑀^levels, but the relation between 𝑃_𝑁_ and 𝑅𝑆𝑁𝑅^a^is assumed to be similar for all a ∈ {*A*, *B*, *C*, *D*, 𝑀}. The four examples from A are highlighted as blue crosses. A polynomial of degree 5 was fitted to the data (red line). (C) Histogram of RNSR over time for the multi-antenna signal (𝑅𝑆𝑁𝑅^𝑀^, blue line), the best single-antenna signal (𝑅𝑆𝑁𝑅^∗^, purple line), all other single-antenna signals (grey lines), and the volumetric antenna signal (red line). The mean (over the measurement period of 2 x 14 h) of 𝑅𝑆𝑁𝑅^𝑀^(dashed blue line) is 6.5 dB larger than the mean of 𝑅𝑆𝑁𝑅^∗^(dashed purple line) and 9.0 dB larger than the mean RNSR of the volumetric antenna (dashed red line). (D) Multi-antenna demodulation is significantly better than single-antenna demodulation for the best single antenna and for the volumetric antenna, as demonstrated by the significantly longer reception periods above a given signal-to-noise ratio. The time below the critical RSNR of 13 dB (fading signal loss) is reduced to 0 %.

In theory, the noise power (variance) of *u*^M^(*t*) is four times larger than that of a baseband signal *u*^a^(*t*), assuming independence of radio amplifier noises. The signal power of *u*^M^(*t*) is 16 times larger than that of a single antenna signal *u*^a^(*t*) when the latter are phase-aligned. Taken together, we expect a 6 dB increase of the 𝑅𝑆𝑁𝑅 for four-antenna demodulation.

By analyzing recorded data of a zebra finch pair over a full day (2 x 14 h measurement period), we found that 𝑅𝑆𝑁𝑅^M^was about 6.5 dB higher on average than the signal-to-noise of the best single antenna (𝑅𝑆𝑁𝑅^∗^, Figure 5C). The total fraction of time during which the 𝑅𝑆𝑁𝑅 was critically low (<13 dB, our operational definition of signal fading) using the multi-antenna demodulation was 0%, compared to 0.62% for the best single-antenna signal (Figure 5D). Thus, our diversity combining approach is very effective.

To verify that simple whip antennas (a form of monopole antenna consisting of a telescopic rod) that we use are not worse than more complex antennas, we recorded a second test dataset of a zebra finch pair (again, 2 x 14 h measurement period) with a single volumetric antenna using single-antenna demodulation. The performance of the volumetric antenna was worse than that of the best simple antenna (Figure 5C, D).

We repeated the 𝑅𝑆𝑁𝑅 analysis on four additional bird groups, yielding an average 𝑅𝑆𝑁𝑅 improvement for multi-antenna demodulation of 6.5 dB (average across 𝑛 = 5 bird groups, group-average range 5.4 – 7.7 dB, Table 1) and a critically low 𝑅𝑆𝑁𝑅 in only 0.00 % (range 0.00 - 0.01%) of the time, compared to 2.71% (range 0.63 – 4.41%, 𝑛 = 5 pairs) of the time for the best single antenna. Thus, our diversity combining robustly tracks frequencies even when the signal vanishes in one or several antennas.

### Vocal segmentation analysis

Microphone and transmitter signals were affected by different failure modes. On microphone channels, some vocalizations were masked by vocal signals of other birds (as e.g., in Figure 6A) or by mechanical noises (Examples B1/B2 in Figure 6B). On transmitter channels, vocal signals reached up to 7 kHz in optimal cases but only up to 1 kHz in the worst case (Figure 6A).

**Figure 6:**
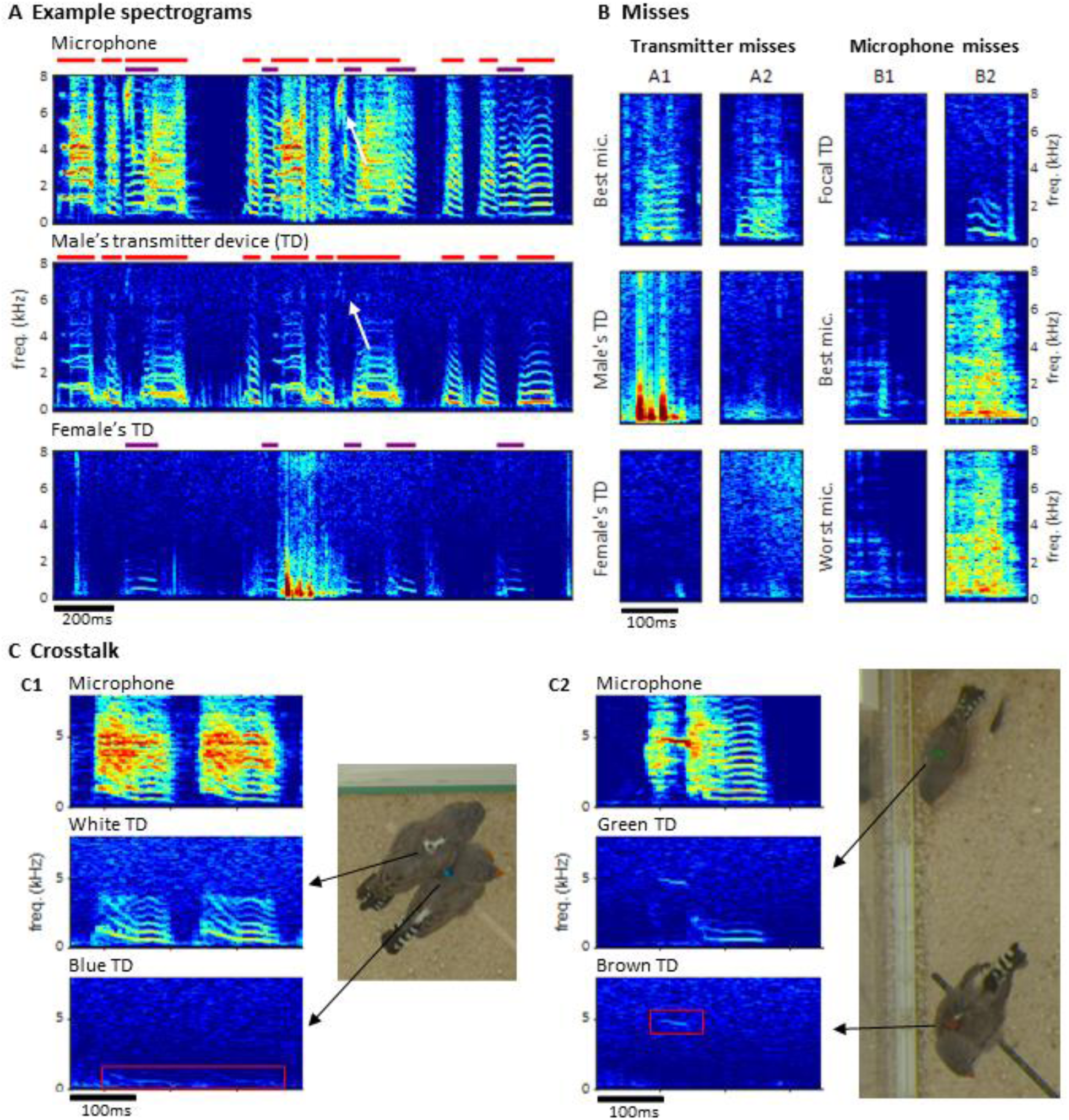
Vocal signals and their segmentation. (**A**) Example spectrograms of vocalizations produced by a mixed-sex zebra finch pair. The songs and calls overlap on the sound spectrogram of a microphone channel (Mic1, top) but appear well separated on acceleration spectrograms of the male’s transmitter channel (middle) and the female’s transmitter channel (bottom). Distinct transmitter-based vocal segments are indicated by red (male) and purple (female) horizontal bars on top of the spectrograms. High-frequency vibrations appear attenuated, but even a high- pitched 7 kHz song note (white arrow) by the male is still visible. (**B**) Example vocalizations that are not visible in the transmitter channels (A1: syllable masked by wing flaps, A2: faint signal) or not visible on either some channels (B1: faint syllable masked by noise) or all microphone channels (B2: syllable masked by loud noise). (**C**) Crosstalk between transmitter channels can occur when birds physically touch (C1), or are near each other (C2). The red rectangles mark the regions in the spectrogram where the vocalization of the focal bird (middle row) leaks into the channel of the non-vocal bird (bottom row).

**Figure 7:**
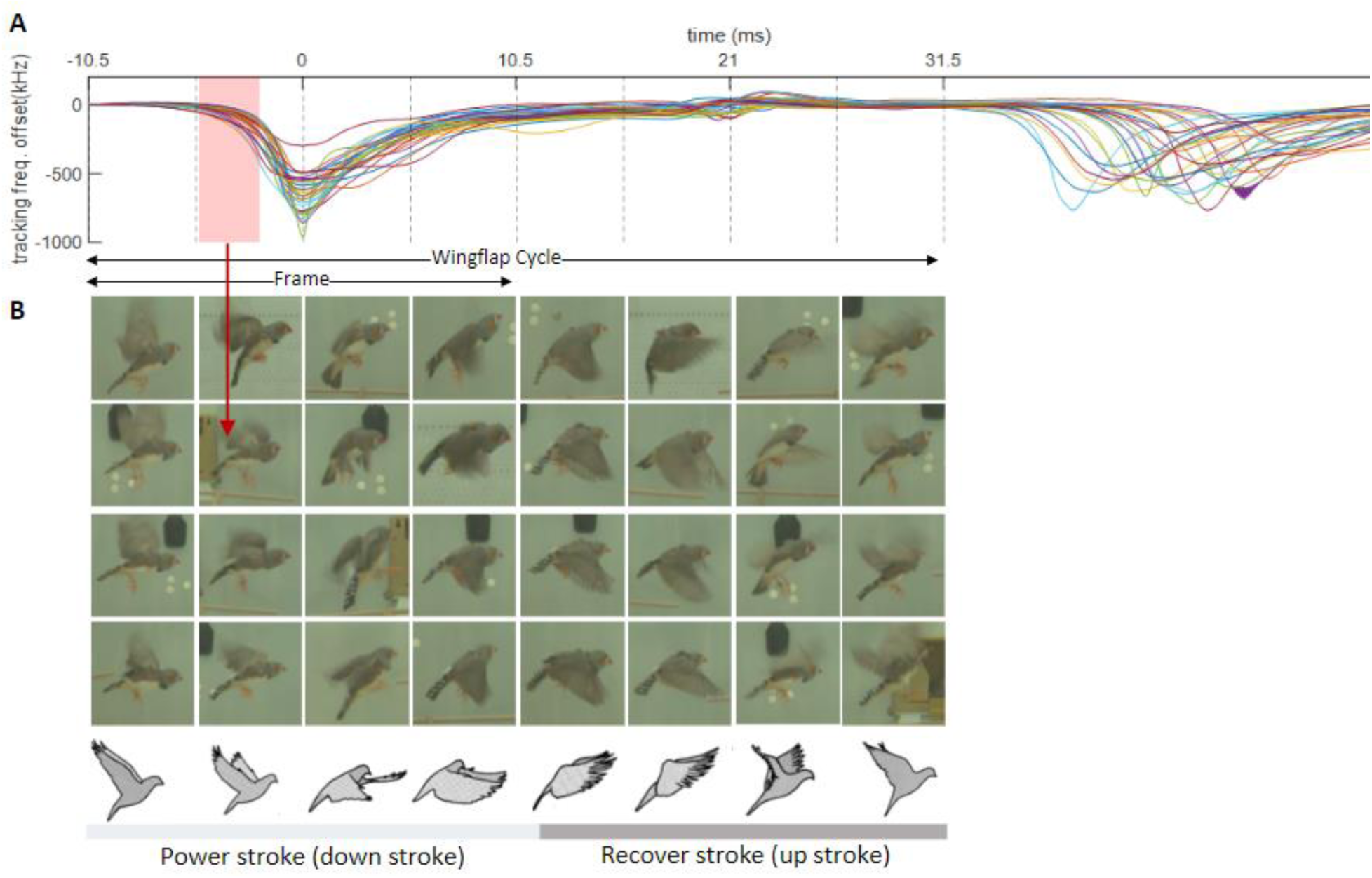
**Depiction of zebra finches’ wing flap movements using transmitter-based time stamping**. Following Brown’s observation of asymmetric hovering in pigeons *(Brown 1963)*, we categorize the wing flap cycle into 8 phases, each represented by four randomly chosen examples (columns). (**A**) The examples’ phase bins we extracted from the time lag between the camera’s exposure window (red shading and dashed lines in A) and the dips in (event-onset subtracted) transmitter signals (n=28 wing-flap events, see Methods). Traces are aligned to the dip, serving as time origin for frame alignment. (**B**) Random video frames (n=32) horizontally arranged in terms of their time lag, illustrating the wing flap movement.

Furthermore, on the transmitter spectrogram, some vocalizations were invisible (Example A2 in Figure 6B), others were masked by simultaneous body movements such as wing flaps or hops (Example A1 in Figure 6B). To quantify these “misses” relative to detected vocalizations in the multimodal data stream, we segmented vocal activity by visual inspection of sound and acceleration spectrograms from microphone and transmitter channels (Supplementary Table S1, see Methods for details). We evaluated the number of missed vocalizations on a given channel relative to a consolidated segmentation that was based on visual inspection of all transmitter and microphone channels combined. We were particularly interested in missed vocalizations that are invisible on a given channel but visible on another channel.

Compared to the miss rate of 1.8% (range 0.1% - 4.6%, 𝑛 = 4 bird groups) associated with a single microphone channel, all microphone channels combined produced a reduction in miss rate by a factor of about 10, which reveals a large benefit of the multi-microphone approach. The microphone misses were mostly very short nest calls such as Example B2 in Figure 6B. The percentage of missed vocalizations on a given transmitter channel of 3.6% (1.8% - 5.8%, 𝑛 = 4 bird groups) was about twice as large as on a single microphone channel. Interestingly, neither the percentage of transmitter nor microphone misses increased with the number of birds in BirdPark. The percentage of microphone misses even decreased in larger bird groups (𝑛 = 4 and 𝑛 = 8 birds); however, this can be attributed to them producing fewer short nest calls, possibly due to the presence of juveniles. Note that the microphone and accelerometer miss rates are not directly comparable because the former stems from two birds and the latter from a single bird. A very small average percentage of 0.1% (range 0.0%-0.5%, 𝑛 = 4 bird groups) of vocalizations were missed on all microphone channels, these vocalizations were all calls masked by loud noises.

The benefit of the transmitter devices is most apparent when vocalizations overlap. We found that 24.1% (range 3.16%-41.0%, 𝑛 = 4 bird groups) of vocalizations overlapped with vocalizations in another bird. The percentage of overlapping vocalizations steeply increased with the number of birds in BirdPark. While microphone channels picked up mixtures of vocal signals (Figure 6A), transmitters did not in most cases. In total, 1.23% of vocalizations were visible on the transmitter of a non-focal bird. Such transmitter channel crosstalk mostly occurred when birds physically touched each other (Example C1 in Figure 6C) but also during loud vocalizations when birds did not touch (Example C2 in Figure 6C). In each instance, the assignment of the vocalization to the vocalizing animal was visually straightforward. In summary, the frequent vocal overlaps in large bird groups prevented the comprehensive analysis of vocalizations from microphone data alone but was possible using the animal-borne sensors. The more birds in the BirdPark, the larger the benefit of the transmitter devices.

### Wing flapping behavior

Our synchronized multimodal recordings, particularly through the combination of video and accelerometer signals, enhance the precision and depth of analyzing movements such as wing flapping. As proof of concept, we demonstrate that the densely sampled transmitter signals can serve as video time stamps that are precisely aligned with the phases of the wing flap cycle, a fundamental movement in birds.

The wing flapping frequency of zebra finches ranges between 26 and 30 Hz (Ellerby and Askew 2007). Our video frame rate of 48 Hz was insufficient to capture such nuanced motion accurately since it is faster than the video Nyquist frequency. Nevertheless, thanks to a randomized sampling method combining the accelerometer signal and the video frames, we were able to comprehensively image the wing flap cycle. Leaning on the transmitters’ frequency shifts caused by body movements, especially the unique shift patterns during wing flapping, we extracted from the transmitter signal the timestamp of an images’ phase in the wing flap cycle. By sorting out the camera’s exposure window, which is considerably shorter than the video frame period, we were able to accurately assign to an image the proper wing flap phase within eight bins (Figure 7, see Methods), i.e., at four times the temporal resolution provided by the camera’s roughly two frames per cycle. Thus, synchronized multi-modal recordings allow for the reconstruction of rapid behaviors and they offer the potential for profound insights into fast-paced social interactions. The limiting factor of behaviors that can be resolved is not set by the camera’s frame rate, but by the shutter speed, i.e., the available light intensity.

## Discussion

By combining transmitter and microphone channels, we showed that vocalization miss rates can be substantially reduced. Currently, there is no reference for miss rates, as to our knowledge, these have not been quantified in previous studies of vocal communication in songbird groups (Ter Maat et al. 2014; Gill et al. 2016; 2015). Being an important parameter, ideally, the miss rate should be evaluated relative to ground-truth, i.e., the true segmentation of vocalizations and background noise. Yet, such a ground-truth data set does not (yet) exist. For example, to perfectly measure the vocal output of a bird would require simultaneous measurements of syringeal labia and muscle activity, of sub-syringeal air pressure, and of tracheal air flow (Goller and Suthers 1996), which are measurements that have not been performed simultaneously in freely moving group-housed birds. Without such a data set or at least an approximation thereof, quantifications of miss rates will be biased towards zero. On the question about acceptable miss rates, the maximally admissible miss rate will depend on the specific research question to be examined, which is why future studies will have to revisit this important parameter.

It remains open how well vocalizations can be separated from the five stationary microphones alone. In groups of mice and using microphone arrays and a camera, it has been shown that vocalizations can be assigned to individual animals by combining sound source localization and animal position tracking (Oliveira-Stahl et al. 2023). But because of frequent vocal jamming in songbirds, we doubt these approaches will make the use of animal-borne transmitters obsolete in song-learning studies to be performed in the BirdPark where subtle song changes need to be tracked in developmental time. Our datasets will be useful for the development and evaluation of such methods, as the transmitter and video channels yield well-resolved information at the individual level.

## Conclusions

We designed and validated a behavioral recording system for up to 8 songbirds, yielding synchronized multimodal data streams. Our custom diversity combining multi-antenna demodulation technique increases the RNSR by 6.5 dB and reduces fading compared with single-antenna demodulation. The wireless devices transmit well-separated vocalizations unless these are either masked by large body movements such as wing flaps, elicited by rare cases of crosstalk, suppressed by poor mechanical coupling between the bird and the device, or lost due to insufficient PLL tracking (which we were unable to eliminate).

While in isolated, non-naturalistic settings, the detection and annotation of vocal segments on a single microphone channel is relatively straightforward, it is much more difficult in social settings and naturalistic environments. We segmented vocal activity by visual inspection of sound and acceleration spectrograms. With respect to these multimodal observations, we find that no sensor channel by itself achieves a miss rate of less than 1%, which is a large number given that we took our measurements in a relatively small, acoustically well isolated environment. Since we might have missed some vocalizations even considering all the channels used, our estimated single-channel miss rates constitute lower bounds, implying that the true miss rate of vocalizations must be higher than what we measured.

To enable automatic segmentation of longitudinal BirdPark data, new methods will need to be developed. While there are many methods for single channel segmentation available (Cohen et al. 2022; Lorenz et al. 2022; Steinfath et al. 2021), only few combine information from multiple sensor channels (Steinfath et al. 2021) and none has made use of accelerometer data so far.

Our system could promote research on the meaning of vocalizations during social behaviors. Of key interest are courtship and reproductive behaviors that have been thoroughly studied (Morris 1954; Caryl 1976; Ullrich, Norton, and Scharff 2016; Morris 1958) albeit often without examining the roles of vocal interactions during mating and pair bonding. Similarly, multimodal group-level studies in the BirdPark could help us better understand the learning strategies young birds use while they modify their immature vocalizations to match a sensory target provided by a tutor. Much of our knowledge on song learning stems from research of isolated animals in impoverished environments (Kollmorgen, Hahnloser, and Mante 2020; Lipkind et al. 2017; Tchernichovski et al. 2001), leaving many questions open about the roles of social interactions (Chen, Matheson, and Sakata 2016; Carouso-Peck and Goldstein 2019)during this process, including by non-singing adult females (Takahashi, Liao, and Ghazanfar 2017).

## Acknowledgements

We thank Aymeric Nager for his help with the design and construction of the BirdPark and Moritz Wohlhauser for creating the illustration in Figure 2D.

## Notes

https://doi.org/10.5281/zenodo.7105196

### Competing Interest Statement

The authors have declared no competing interest.

### Summary of Updates

Author M.D.Rocha is removed from the author list. Text is modified in session 'Animals and experiments'

