## Supplemental Table 1 for "Multimodal system for recording individual-level behaviors in songbird groups"

### Supplementary Table S1

| # | Experiment | copExpBP08 | copExpBP09 | juvExpBP01 | juvExpBP03 | Total |
|---|---|---|---|---|---|---|
| **2** | **Segmented data** | 1 x 7 mins | 1 x 7 mins | 7 x 1 min | 7 x 1 min | **28 min** |
| **3** | **# birds** | 2 | 2 | 4 | 8 | **16** |
| **4** | **# vocalizations** | 537 | 570 | 709 | 861 | **2677** |
| **5** | **# voc. missed on tr. channel** | 31 (5.8%) | 10 (1.8%) | 37 (5.2%) | 18 (2.1%) | **96 (3.6%)** |
| **6** | **# voc. missed on all mic. channels** | 1 (0.2%) | 3 (0.5%) | 0 (0.0%) | 0 (0.0%) | **4 (0.2%)** |
| **7** | **# voc. missed on Mic1 channel** | 20 (3.7%) | 26 (4.6%) | 1 (0.1%) | 2 (0.2%) | **49 (1.3%)** |
| **8** | **# voc. unassigned** | 0 (0.0%) | 0 (0.0%) | 0 (0.0%) | 1 (0.0%) | **1 (0.0%)** |
| **9** | **# voc. with overlaps** | 18 (3.4%) | 18 (3.2%) | 257 (36.2%) | 353 (41%) | **646 (24.1%)** |
| **10** | **# voc. with crosstalk** | 0 (0.0%) | 0 (0.0%) | 30 (4.2%) | 3 (0.4%) | **33 (1.2%)** |
| **11** | **# uncertain segments** | 3 | 3 | 0 | 0 | **6** |

Table S1: Statistics of missed vocalizations on a transmitter channel (5th row, cf. Example A1 and A2 in Figure 6B), on all microphone channels (6th row, cf. Example B2 in Figure 6B), and on an individual microphone channel (Mic1, 7th row, cf. Example B1 in Figure 6B). We report the number of vocalizations for which an assignment to a bird was not possible (8th row); the number of vocalizations that overlap with at least one vocalization of another bird (9th row, cf. example in Figure 6A); the number of vocalizations with transmitter channel crosstalk (10th row, cf. Examples in Figure 6C); and the number of uncertain segments (11th row). See Methods for details.
